# Interrogating the *Escherichia coli* epitranscriptome via CRISPR interference and Nanopore native RNA sequencing

**DOI:** 10.64898/2026.04.13.718120

**Authors:** Miranda E. Pitt, Jianshu Zhang, An N. T. Nguyen, Michael B. Hall, Leila Jebeli, Leo A. Featherstone, Garry S.A. Myers, Nichollas E. Scott, Lachlan J.M. Coin

**Author notes:** Corresponding authors: Miranda E. Pitt, and Lachlan J.M. Coin. These authors contributed equally (Miranda E. Pitt, Jianshu Zhang).

## Abstract

Epitranscriptomics has recently gained significant momentum due to technological advances and translational applications, however, studies on bacterial RNA modifications remain limited. Bacterial RNA remains notoriously prone to degradation and methodologies to investigate the epitranscriptome are challenging. Prior research has shown RNA modifications modulate antimicrobial resistance, virulence and pathogenicity.

This research employed CRISPR interference to knock down five known *Escherichia coli* rRNA modification genes (*rlmF*, *rlmJ*, *rluD*, *rsmF* and *rsmG*) in three *E. coli* strains. These isolates underwent growth curves, proteome analysis and native RNA sequencing CRISPRi adequately silenced the majority of RNA modification genes in *E. coli* (>80% reduction). Significant growth delays were associated with *rlmF*, *rsmF* and *rsmG* repression. Unique protein pathways corresponding with RNA modification loss were found for *rlmJ* (TreB, XylF), *rluD* (CysH, HycB, PutP, TrpB), *rsmF* (EvgA) and *rsmG* (OppC). Known rRNA modification sites for *rluD* (Ψ) and *rsmG* (m7G) were detected from analysis of nanopore electrical signal, however, only a weak signal was apparent for m6A (*rlmF, rlmJ*) and m5C (*rsmF*) modifications. The inhibition of rRNA modifications resulted in mRNA modification changes including for genes *ompC*, *cspC*, *dbhA*, *dbhB* and *secY*.

Our work provides an approach for unravelling the epitranscriptome of *E. coli* and gain insight into its functional role.

## INTRODUCTION

RNA facilitates various cellular processes including coding, decoding, regulation and gene expression. This single stranded molecule comprises of a sugar (ribose)-phosphate backbone and a combination of four nitrogenous bases (adenine (A), cytosine (C), guanine (G) and uracil (U)). There are three major types of RNAs including: ribosomal RNA (rRNA) that forms the ribosomes; transfer RNA (tRNA) which transport amino acids to the ribosomes for protein synthesis and messenger RNA (mRNA) which is converted to proteins via the ribosomes. Following transcription, RNA bases can be chemically altered which influence RNA stability and function. The full set of RNA modifications is commonly referred to as the epitranscriptome [1].

To date, >170 types of RNA modifications have been identified [2, 3], however, this has commonly been restricted to rRNA and tRNA modifications. This is primarily due to rRNA and tRNA occurring in higher abundance (rRNA: ∼80-90%, ∼10-15% tRNA and ∼1-5% mRNA) [4]. Methods for detecting RNA modifications remain laborious, requiring high purity and concentration of the RNA encoded gene of interest. Furthermore, restricting the detection of differing modifications and multiple RNA types being interrogated simultaneously [1,5]. Recent advances in sequencing technologies may overcome these shortcomings [6]. In 2017, Oxford Nanopore Technologies (ONT) released a sequencing technique known as direct or native RNA sequencing. Other sequencing methods rely on the conversion of RNA to cDNA which removes RNA modifications and converts the U base to T in the process [6,7].

Most published work utilising direct RNA sequencing has been performed on human, yeast and viral RNA [7–13]. These studies are restricted to a subset of commonly occurring modifications (N6-methyladenosine (m6A) [7–9], 5-methylcytosine (m5C) [7], A to inosine editing (A-to-I) [8]. Previous bacterial studies using ONT direct RNA sequencing have primarily focused on *E. coli* rRNA. This included detecting known 7-methylguanosine (m7G), 2’-O-methyl and pseudouridine (Ψ) modifications on 16S rRNA [14,15]. Notably, one study interrogated all known *E. coli* rRNA (16S and 23S) sites using direct RNA sequencing which encompassed 36 sites and 17 modifications [16]. As a result, there is still limited information on the modifications occurring in bacterial mRNA. Despite this, emerging research is indicating that various modifications are present and play important roles. Similar to other organisms, modifications such as m6A [17], m5C [18], A-to-I [19], m7G [14] and Ψ [15] have been identified in bacteria. Several studies have shown that by removing the genes enabling tRNA modifications can impede on its ability to infect, grow and cause disease in a host [20–22]. Modifications to rRNAs enable the evasion of antibiotics including aminoglycoside, chloramphenicol and macrolide antibiotic classes [23,24]. Recently, the loss of several nonessential tRNA and rRNA modifications were identified to facilitate tolerance to sub-inhibitory concentrations for various antibiotics in *Vibrio cholerae* [25].

The key challenges entailed in investigating the bacterial epitranscriptome is the rapid degradation of RNA, with most transcripts estimated to have <1-minute half-lives [26]. Hence, utilising shorter RNA preparation times compared to laborious protocols such as immunoprecipitation and chemical treatments (e.g. bisulfite) are favourable [6]. Direct RNA sequencing provides quicker turnaround times, however, due to structural differences with bacterial RNA (lack of polyadenylation), this sequencing chemistry was not compatible [27]. Previously, our research has utilised artificially adding poly(A) tails to bacterial transcripts to enable compatibility with direct RNA sequencing [27, 28].

In this study, we have implemented CRISPR interference (CRISPRi) to target five bacterial rRNA modification genes: Ribosomal RNA large subunit methyltransferase-F (*rlmF*),-J (*rlmJ*), Ribosomal RNA small subunit methyltransferase-F (*rsmF*),-G (*rsmG*) and Ribosomal large subunit pseudouridine synthase D (*rluD*). As only one A-to-I RNA modification is known in bacteria (*tadA*, tRNA [19]), an additional gene Adenosine deaminase (*add*) was explored to see if there was any potential RNA modification activity.

CRISPRi differs from CRISPR as it will knock down (KD), rather than knock out (KO), genes and has been previously applied to bacteria [29–32]. Dissimilar to CRISPR approaches, the Cas9 endonuclease (DNA cutting) activity is inactivated in CRISPRi which harbours a deactivated Cas9 (dCas9) [29]. Hence, a guide RNA (gRNA) can be designed to target the gene of interest, dCas9 will bind to this DNA/gRNA complex and block the transcription of this gene. Identifying the potential functional role of RNA modifications in bacteria remains elusive. In order to initially detect these RNA modifications, direct RNA sequencing was used to interrogate all RNA types including mRNA. To determine the impact of RNA modification loss, *E. coli* KD isolates underwent growth curves to assess fitness and downstream influences on proteins via proteomics.

## MATERIALS AND METHODS

### CRISPRi design

The pFD152 plasmid was a gift from David Bikard (Addgene plasmid #125546; http://n2t.net/addgene:125546; RRID:Addgene_125546). The pFD152 plasmid (Addgene [Watertown, MA, USA]) encodes an inducible dCas9 (via anhydrotetracycline (aTc)), an antibiotic selectable marker for spectinomycin and a restriction site for gRNA insertion using BsaI [29]. The six target genes were cross referenced on the RegulonDB browser (https://regulondb.ccg.unam.mx/) and only genes not in operons or were the last gene of an operon were selected [33]. Sequences were extracted (https://www.bioinformatics.org/sms2/range_extract_dna.html) using annotations and assemblies provided by the ATCC Genome portal (±200bp from start site). CRISPRi constructs were designed via the CRISPR@Pasteur browser (https://crispr-browser.pasteur.cloud/guide-rna-design) which provided a score for off-target sites in the genome [34]. gRNAs (Merck) were selected based on positions close to promotor and start of genes with the least off-target sites (Supplementary Table S1).

### Bacterial cultures and growth conditions

*E. coli* strains (ATCC 11775, 8739, 25922, BAA-2452) were sourced from the American Type Culture Collection (VA, USA). *E. coli* strains were selected based on complete genome availability through the ATCC Genome Portal (https://genomes.atcc.org). When selecting strains suitable for CRISPRi experiments, genomes were queried across ResFinder v4.6.0 (90% identity, 60% minimum length) [35] to ensure no resistance to selectable markers spectinomycin and tetracyclines and PlasmidFinder v2.1 (95% minimum identity, 60% minimum coverage) [36] to negate replicons which may compete with the CRISPRi plasmid (pFD152). *E. coli* strains were stored in Lysogeny Broth (LB) with 20% (v/v) glycerol at-80°C. When required for subsequent assays, glycerol stocks were struck out on LB agar (overnight, 37°C), inoculated in LB and grown overnight at 37°C, 180 rpm.

### CRISPRi transformations

The gRNA cassette was introduced into the pFD152 plasmid as previously described [29]. Minor modifications to the protocol include using an optimised BsaI-HF®v2 (New England Biolabs) to insert the gRNA into pFD152 which was transformed into PIR2 cells (Thermo Fisher Scientific) rather than DH5α and positive colonies were selected on 50 µg/ml spectinomycin LB agar plates. Plasmids were extracted using the QIAprep Spin Miniprep kit (Qiagen). To confirm the plasmid presence and gRNA insertion, regions flanking externally and internally of the gRNA (Supplementary Table S2) were amplified using Hot Start Taq 2X Master Mix (New England Biolabs) using cycling conditions: 35X of 95°C 20 seconds, 55°C 30 seconds and 68°C 1 minute. To ensure no errors had arisen in the plasmids, the pFD152 external primers along with Phusion™ High-Fidelity DNA Polymerases (2 U/μL) (Thermo Fisher Scientific) were amplified under the following cycling conditions: 35X of 98°C 10 seconds, 55°C 30 seconds and 72°C 30 seconds. PCR products were purified using the GeneJET PCR Purification Kit (Thermo Fisher Scientific) and amplicon identity confirmed via Sanger sequencing (AGRF, Victoria).

*E. coli* ATCC electrocompetent cells were prepared as previously described [37] with a minor change: centrifugation steps at 6,000xg. *E. coli* isolates were transformed using electroporation (2.5 kV, 5.3±0.2 ms) with 0.2 µm gap cuvettes and a MicroPulser electroporator (Bio-Rad Laboratories). Positive colonies were selected on 50 µg/ml spectinomycin (Merck) LB agar plates. Plasmids were extracted using the QIAprep Spin Miniprep kit (Qiagen) and validated using external and internal primers (Supplementary Table S2) with the Hot Start Taq 2X Master Mix (New England Biolabs) as described above.

### RNA extractions and DNA depletions

RNA extractions and DNA depletions were conducted as previously described with minor changes [27]. Briefly, the overnight culture was sub-cultured (1:100 dilution) into LB and grown to mid-log phase (OD600 = 0.5-0.6). For isolates containing the pFD152 plasmid, overnight cultures and sub-cultures were supplemented with 60 µg/ml spectinomycin hydrochloride (78.6±5.3 % purity, Merck). For KD cultures, only sub-cultures contained additional 0.5 µg/ml aTc hydrochloride (≥95.0 % purity, Merck) with 60 µg/ml spectinomycin and cultures were protected from light exposure. RNA was extracted using the PureLink^TM^ RNA Mini Kit (Thermo Fisher Scientific) with Homogenizer columns (Thermo Fisher Scientific). DNA was removed using the TURBO DNA-free^TM^ kit (Thermo Fisher Scientific) with 4 U TURBO DNase incubated at 37°C for 30 minutes. RNA was purified with RNAClean XP beads (1.8X beads: RNA (v/v)) (Beckman Coulter).

### Quantity control

RNA (RNA High Sensitivity kit) and DNA (1X dsDNA kit) was quantitated using a Qubit®4.0 Fluorometer (Thermo Fisher Scientific). Purity was determined via a NanoDrop 2000 Spectrophotometer (Thermo Fisher Scientific). RNA fragmentation was determined using an Agilent High Sensitivity RNA ScreenTape on a 4200 TapeStation (Agilent Technologies).

### qRT-PCR knock down validation

RNA (1 µg total RNA (DNA-depleted)) underwent reverse transcription using SuperScript III (Thermo Fisher Scientific) and random primers from the High-Capacity cDNA reverse transcription kit (Thermo Fisher Scientific) under conditions: 65°C 5 minutes, 25°C 10 minutes, 37°C 30 minutes, 50°C 30 minutes and 70°C 15 minutes. Genes of interest were amplified via the SYBR Select Master Mix (Thermo Fisher Scientific) under the following conditions: 50°C 2 minutes, 95°C 2 minutes followed by 50X 95°C 15 seconds and 60°C 1 minute followed by melt curves. qRT-PCR was run on a QuantStudio Flex 7 PCR system (Thermo Fisher Scientific) and analysis with the QuantStudioTM Real-Time PCR Software. Primers used are displayed in Supplementary Table S2 and were normalized to housekeeping genes *rssA* and *rpoS*. Triplicates were averaged and expression determined via the 2^-ΔΔCT^ method [27]. Significance was defined as p<0.05 via a Mann-Whitney test.

### Growth curves

Growth curves were conducted as previously described [38]. Overnight cultures grown in 60 µg/ml spectinomycin were diluted to OD600 = 0.1. 150µl of diluted culture was transferred to 96-well Clear Flat Bottom Polystyrene Not Treated Microplates (Corning) (n≥3 technical replicates, n=4 biological replicates). Control isolates were supplemented with 60 µg/ml spectinomycin and KD strains were additionally supplemented with 0.5 µg/ml aTc. A breathe-easy membrane (Merck) sealed the plate, growth was read on a CLARIOstar Plus Microplate reader (BMG Labtech) for 24 hours (readings every 1 hour) with continuous shaking (200 rpm). Doubling time was calculated as follows:

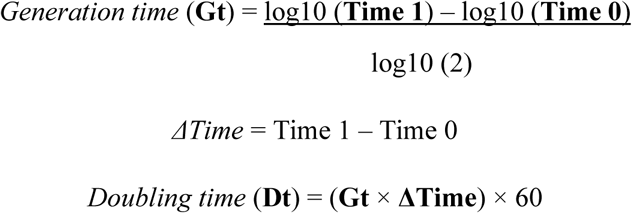

Time 0 is the start and Time 1 is the end of exponential phase. Significance was determined using a two-tailed Welch’s t-test.

### Sample preparation for Proteomic analysis

Control (grown in 60 µg/ml spectinomycin) and KD (60 µg/ml spectinomycin, 0.5 µg/ml aTc) isolates (n=4 biological replicates) were prepared for protein extractions. Briefly, overnight cultures (16 hours) were diluted to OD600 = 1.0, centrifuged at 6,000xg 5 minutes 4°C and washed with ice-cold PBS. Frozen whole bacterial pellets were prepared using the in-StageTip preparation approach as previously described [39]. Cells were resuspended in 4% sodium deoxycholate (SDC), 100 mM Tris pH 8.0 and boiled at 95°C with shaking for 10 minutes to aid solubilisation. Samples were allowed to cool for 10 minutes and then boiled for a further 10 minutes before the protein concentrations were determined by bicinchoninic acid assays (Thermo Fisher Scientific). 100μg of protein for each biological replicate were reduced / alkylated with the addition of Tris-2-carboxyethyl phosphine hydrochloride and iodoacetamide (final concentration 20 mM and 60mM respectively), by incubating in the dark for 1 hour at 45°C. Following reduction / alkylation samples were digested overnight with Trypsin (1/33 w/w Solu-trypsin, Sigma) at 37°C with shaking at 1000rpm. Digests were then halted by the addition of isopropanol and TFA (50% and 1% respectively) and samples cleaned up using home-made SDB-RPS StageTips prepared according to previously described protocols [39–41]. SDB-RPS StageTips were placed in a Spin96 tip holder 2 to enable batch-based spinning of samples and tips conditioned with 100% acetonitrile; followed by 30% methanol, 1% TFA followed by 90% isopropanol, 1% TFA with each wash spun through the column at 1000 x g for 3 minutes. Acidified isopropanol / peptide mixtures were loaded onto the SDB-RPS columns and spun through, and tips washed with 90% isopropanol, 1% TFA followed by 1% TFA in Milli-Q water. Peptide samples were eluted with 80% acetonitrile, 5% ammonium hydroxide and dried by vacuum centrifugation at room temperature before being stored at-20°C.

### Reverse phase Liquid chromatography–mass spectrometry

Prepared digested proteome samples were re-suspended in Buffer A* (2% acetonitrile, 0.01% trifluoroacetic acid) and separated using a two-column chromatography setup composed of a PepMap100 C_18_ 20-mm by 75-µm trap and a PepMap C_18_ 500-mm by 75-µm analytical column (Thermo Fisher Scientific). Samples were concentrated onto the trap column at 5 µl/min for 5 min with Buffer A (0.1% formic acid, 2% DMSO) and then infused into a Orbitrap Q-Exactive plus at 300 nl/minute via the analytical columns using a Dionex Ultimate 3000 UPLCs (Thermo Fisher Scientific). 90-minute analytical runs were undertaken by altering the buffer composition from 2% Buffer B (0.1% formic acid, 77.9% acetonitrile, 2% DMSO) to 22% B over 60 min, then from 22% B to 40% B over 10 min, then from 40% B to 80% B over 5 min. The composition was held at 80% B for 5 min, and then dropped to 2% B over 2 min before being held at 2% B for another 8 min. The Q-Exactive plus Mass Spectrometer was operated in a data-dependent mode switching between the collection of a single Orbitrap MS scan (375-1400 m/z and a resolution of 70k) and up to 15 f MS/MS HCD scans of precursors (Stepped NCE 28,30,42%, with a maximal injection time of 50 ms with the Automatic Gain Control set to 5e^4^ and the resolution to 17.5k).

### Proteomic data analysis

Identification and LFQ analysis were accomplished using MaxQuant (v2.4.7.0) [42] with datasets searched against the *E. coli* 11775 (Uniprot: UP000028628), *E. coli* ATCC 8739 (Uniprot: UP000000317) or *E. coli* ATCC 25922 (Uniprot: UP000028955) proteomes with the addition of the dCas9 protein sequence to allow the monitoring of silencing induction. Searches were undertaken using a trypsin specificity and allowing for oxidation on Methionine as well as carbamidomethyl on Cysteine as a fix modification. The LFQ and “Match Between Run” options were enabled to allow comparison between samples. The resulting data files were processed using Perseus (v1.4.0.6) [43] to compare the growth conditions using student t-tests. For LFQ comparisons biological replicates were grouped and missing values imputed based on the observed total peptide intensities with a range of 0.3σ and a downshift of 1.8σ. To enable comparison between strains, protein groups were mapped to the *E. coli* K12 reference (Uniprot: UP000000625) using the PATRIC proteome comparison tool [44].

### RNA processing for Oxford Nanopore Technologies direct RNA sequencing

RNA extracted from replicates which harboured the highest KD efficiency, determined via qRT-PCR, were selected for sequencing. Two controls include the initial *E. coli* isolate and pFD152 lacking the gRNA (empty vector) with induced expression of dCas9 via 0.5 µg/ml aTc. To determine if the RNA modification had been sufficiently removed from rRNA, total RNA (DNA-depleted) was initially run on a flongle (R9.4.1) flow cell. Total RNA (1 µg) was polyadenylated prior to sequencing library preparation and using the Poly(A) Polymerase Tailing Kit (Astral Scientific) as previously described [27] with minor adjustments: incubated at 37°C for 5 minutes and purified using RNAClean XP beads ((1.8X beads: RNA (v/v)) (Beckman Coulter). RNA poly(A)+ (∼250ng) underwent direct RNA sequencing library preparation (SQK-RNA002). 20ng of library was added onto the flongle flow cell and run on the GridION platform (ONT) for 48 hours.

To detect potential mRNA modifications, total RNA (DNA-depleted) (8 µg) underwent a rRNA depletion as per described [27]. The MICROBExpress^TM^ Bacterial mRNA Enrichment Kit (Thermo Fisher Scientific) was used followed by a TapeStation 4200 run to ensure 16S and 23S rRNA had been significantly depleted (≥85%). Approximately 1 µg of RNA (rRNA-depleted) underwent poly(A) tailing as described above and ∼500ng of RNA poly(A)+ was used for direct RNA sequencing library preparations (SQK-RNA002). Libraries were sequenced on R9.4.1 flow cells on a GridION for 72 hours

### Base-calling, alignment and quantification

All raw fast5 signals that passed the qscore threshold (>7) were basecalled with dorado (v0.5.3) using model rna002_70bps_hac@v3. Fastq files from dorado outputs were used for alignment and quantification. Minimap2 (v2.24) [45] was implemented for alignment using parameter “-y-ax map-ont-L --MD”. Flongle sequencing reads were only mapped to 16S plus 23S rRNA reference and GridION mRNA sequencing reads were aligned to 16S plus 23S rRNA and *Escherichia coli* transcriptomic reference from 3 different strains. Samtools (v1.16.1) [46] were employed to compile the statistics on all sequencing data. Read Qscores were calculated using python package “pysam”. Gene expression on GridION samples were quantified using Salmon (v1.10.1) [47] against transcriptomic reference from 3 strains.

### Unifying annotation of open reading frames between strains

To compare modification sites between 3 different *Escherichia coli* strains, we employed ggcaller (v1.3.0) [48] to standardize the annotation of open reading frames. Based on ggcaller documentation, in default mode, bacterial database from uniprot was employed by DIAMOND [49] to annotate the input reference genomes. The gene names mentioned in results and discussion section were based on ggcaller unified annotation.

### RNA modification site calling

We first used f5c (v1.3) to index the reads with sequence summary files created from basecalling. Then, we event-aligned all signal events with aligned reads using f5c (v1.3) as well under parameters: --scale-events --print-read-name --secondary=yes --min-mapq 0 --samples --rna. We then implemented nanocompore (v2.0.0) “eventalign_collapse” to collapse aligned events into qlite format datasets. To compare events from different conditions (one knockdown vs one control), we used nanocompore (v2.0.0) [50].

### P value combination and plotting

To combine the p value from different strains and controls under the same knockdown, python package scipy(v1.16.1) was used to perform Fisher’s combined probability test. The p value adjustment was calculated using scipy(v1.16.1) as well. Plots were created using R (v4.4.0) with ggplot2 package(v3.5.2). Logo plots for motif analysis were generated using ggseqlogo [51].

## DATA AVAILABILITY

The direct RNA sequencing datasets are available in the NCBI repository BioProject PRJNA1348313 (www.ncbi.nlm.nih.gov/bioproject/PRJNA1348313) to access fastq files. Raw data (fast5) files will be made available via figshare. The mass spectrometry proteomics data has been deposited in the Proteome Xchange Consortium via the PRIDE partner repository. ATCC 11775 can be accessed via the Project accession: PXD069957 and Token: QnzuKbXaAnr4; ATCC 25922 via the Project accession: PXD069976 and Token: zgdpQuTlp3ms; and ATCC 8739 via the Project accession: PXD069998 and Token: xGVt85VY5w0F.

## CODE AVAILABILITY

Code generated for this manuscript is available at github: https://github.com/abcdtree/RNA-mods-paper

## RESULTS

### Expression of RNA modification genes and knock down efficiency

To test the efficiency of the CRISPRi model, four ATCC *E. coli* isolates were selected with differing characteristics. ATCC 11775 harbours a 4.90Mb genome with a 131 kb plasmid (replicon: IncFIB) and no antibiotic resistance. ATCC 25922 has a 5.13Mb genome with four mobile elements: 48.5 kb, 24.2 kb (IncX1), 3.2 kb and 1.9 kb and no antibiotic resistance. ATCC 8739 has a 4.74Mb genome with no plasmids and no antibiotic resistance. ATCC BAA-2452 has a 4.78Mb genome with a 144kb (IncFIA) and 138kb (IncC) plasmid. ATCC BAA-2452 is a multidrug-resistant isolate with resistance to beta-lactams/ quinolones (*blaNDM-1*, *blaCMY-6*), aminoglycosides (*aac(6’)-Ib3*, *rmtC*) and sulphonamides (*sul1*).

The expression of the five known RNA modification genes and *add* were evaluated using qRT-PCR. Under standard growth conditions in LB, all genes were expressed (Supplementary Figure S1). In all four *E. coli* isolates, *add* followed by *rlmJ* had the highest relative expression. All expression trends were the same when normalised to both housekeeping genes *rpoS* and *rssA*. The lowest relative expression across all isolates was observed for *rlmF* and *rsmF*.

Two gRNAs were designed and tested against ATCC 11775 initially (Figure 1A). Only a few PAM sites could be detected within ±200bp of the gene start site as well as avoiding inhibiting other closely neighbouring genes detected by annotations and RegulonDB. Overall, similar levels of repression were observed for each gRNA with the exception of *rlmF* (80.65% for gRNA 1 compared to 34.89% for gRNA 2). The majority of RNA modification genes could be reduced by ∼≥80% however, *rsmG* could only be reduced by ∼54%. Thus, the design for gRNA 1 was used for other strains due to fewer off-target sites (minimum off-target score: ≥0.71 gRNA 1 and ≥0.29 for gRNA 2), exhibiting the highest KD and targeting a more suitable site in the gene (closer to the start site) (Supplementary Table S1).

**Figure 1.**
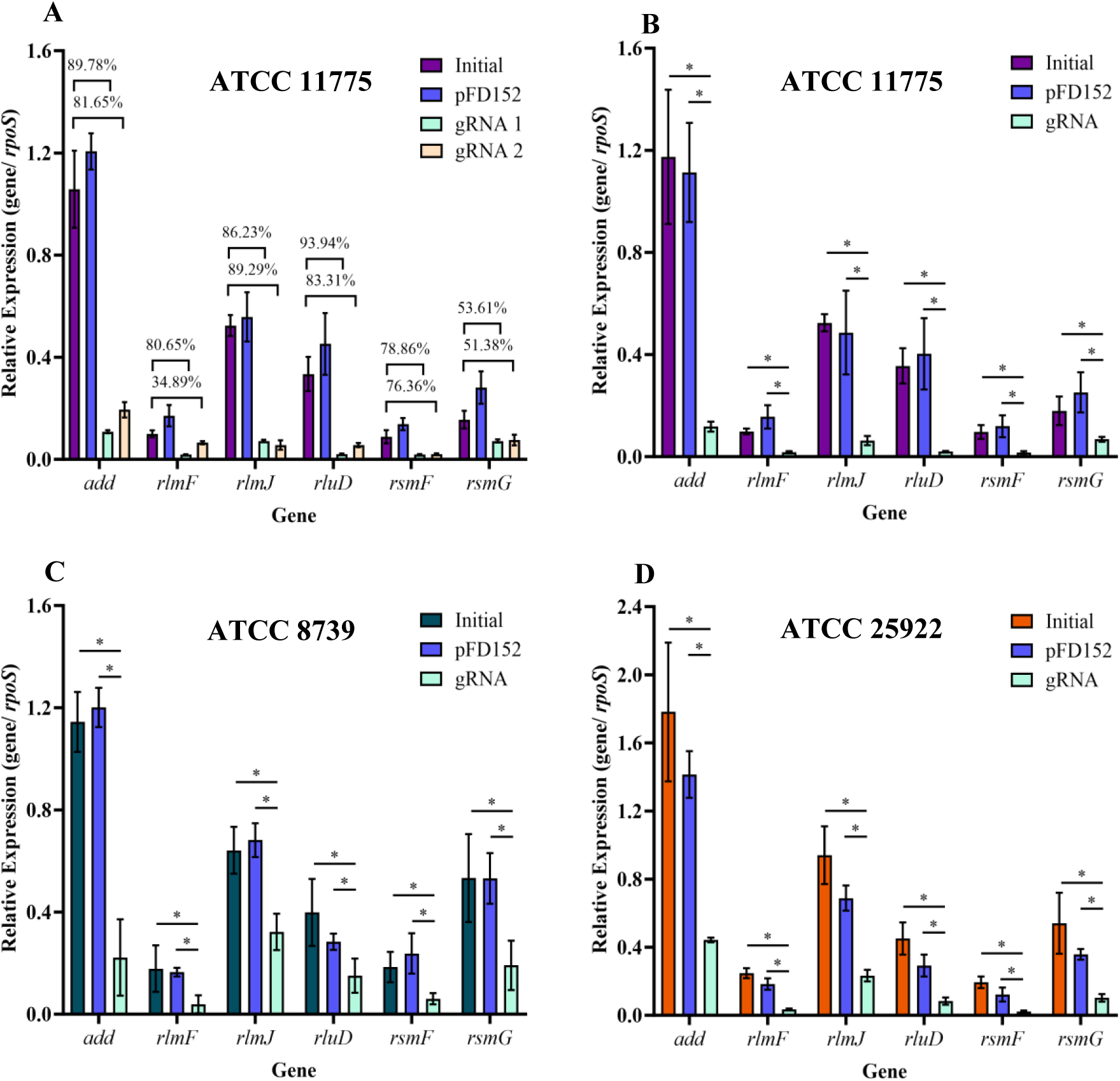
Efficiency of gene knock down via pFD152 in three *E. coli* strains. **A)** ATCC 11775 KD of genes using 2 different gRNAs with control pFD152 empty vector (blue) induced to express dCas9 (n=3±SD). Percentages (%) indicate average reduction in expression between initial and 2 gRNAs. **B)** ATCC 11775 KD for selected gRNA with additional biological replicates (n=4±SD). **C)** ATCC 25922 for selected gRNA (n=4±SD). **D)** ATCC 8739 for selected gRNA (n=4±SD). All normalized to housekeeping gene *rpoS* with significance as p≤0.05 (*) via Mann-Whitney test.

The pFD152 plasmid was further transformed into ATCC 25922 and ATCC 8739. Numerous attempts to transform ATCC BAA-2452 were unsuccessful with adjustments such as increasing pFD152 amount (∼10ng to 50ng), increasing culture volume on transformation plates and incubating plates longer. Whilst there was evidence that pFD152 could be transformed into ATCC BAA-2452, a major fitness cost was observed and due to these difficulties, this strain was not carried forward in this study. For all three *E. coli* isolates (Figure 1B-D), a significant reduction (p<0.05, Mann-Whitney test) was observed when inducing the expression of dCas9 and the gRNA for all six gene targets. No significant difference was seen when comparing the initial isolate and pFD152 empty vector stimulated to express dCas9. Of note, ATCC 25922 had the most variance in its gRNA KD efficiency (Figure 1C). ATCC 25922 transformants struggled to generate high enough pFD152 expression as some cell death was observed when grown in 60 µg/ml spectinomycin. Spectinomycin concentrations were adjusted and a new batch of ATCC 25922 was obtained, however, similar results were observed. Throughout this study, 60 µg/ml spectinomycin was used for ATCC 25922 to ensure the pFD152 plasmid was retained and the cell death (sedimentation) was avoided when samples were required for extractions.

### Impact of CRISPRi on bacterial growth

The activation of dCas9 between ATCC 11775 and ATCC 8739 showed differing impacts on bacterial doubling time (Supplementary Figure S2). In ATCC 11775, a high fitness cost was observed for the empty pFD152 plasmid control once dCas9 was activated (pFD152 (+), dCas9 (+), no gRNA) compared to the initial isolate (pDF152 (-)) (p<0.0001, two-tailed Welch’s t-test) and pFD152 control (pFD152 (+), dCas9 (-), no gRNA) (p<0.0001) (Supplementary Figure S2A). A significant growth reduction was observed between initial control and pFD152 for all six genes repressed (pFD152 (+), dCas9 (+), gRNA (+)) (p<0.0001). A significant difference between initial and pFD152 control was observed (p= 0.0016) but no significant difference between these controls and the non-activated isolates. Compared to the pFD152 (+) (dCas9 (+), no gRNA) control, only a significant difference was observed between *rluD* (p=0.0001) and *rsmF* inhibited (p=0.0012). Of note, this pFD152 control compared to *rlmF* inhibited resulted in a close to significant p value of 0.051. For ATCC 8739, the growth reduction was less apparent when pFD152 was introduced into this strain and with dCas9 activation (Supplementary Figure S2B). The only significant growth delays observed were between pFD152 (+) (dCas9 (+), no gRNA) control and *rlmF* inhibited (p=0.0303). Between gene control (pFD152 (+), dCas9 (-), gRNA (-)) and inhibited (pFD152 (+), dCas9 (+), gRNA (+)), significant growth delays were seen for *rlmF* (p=0.042), *rluD* (p=0.0124) and *rsmF* (p=0.0358).

### Unique protein pathways impacted in *E. coli* CRISPRi isolates

Strains ATCC 11775, 25922 and 8739 underwent protein extraction and analysis with comparisons between KD (pFD152 (+), dCas9 (+), gRNA (+)) and pFD152 gene control (pFD152 (+), dCas9 (-), gRNA (-)). Overall, 2,383 proteins were detected in ATCC 11775, 2,206 in ATCC 25922 and 1,827 in ATCC 8739. As a further control, the pFD152 empty vector and pFD152 dCas9 activated samples were run to determine the influence of dCas9 expression (Supplementary Fig S3). ATCC 11775 showed a significant increase ((fold-change(log2) >2,-log10 (p-value) >2)) of dCas9 (Supplementary Figure S3, Supplementary Figure S4). ATCC 8739 also showed a significant increase in dCas9 except in the *add* samples which was very close to the cut off (dCas9:-1.90, 4.56) and ATCC 25922 was unable to increase dCas9. Only subtle reductions were seen for the KD protein of interest with the only significant changes observed for ATCC 11775 RlmJ and ATCC 8739 RlmJ and RsmG (Supplementary Figure S4). However, near the cut-off were ATCC 11775 RlmF (fold-change(log2):-1.93,-log10 (p-value): 3.18), RluD (-1.55, 3.12); and ATCC 8739 Add (-1.17, 3.52) and RluD (-1.95, 2.48) (Supplementary Figure S4). For ATCC 8739, protein levels for RlmF and RsmF were too low to be detected.

Significant protein abundance changes identified in the pFD152 control (pFD152 (+), dCas9 (+), no gRNA versus pFD152 (+), dCas9 (-), no gRNA) were filtered out from all datasets to refine the pathways impacted by the repression of genes of interest (Supplementary Figure S3G). Several proteins impacted by CRISPRi were detected in at least 2 *E. coli* strains (Table 1). The reduction of Citrate lyase alpha chain (CitF) was observed in *add*, *rlmF, rlmJ* and *rluD* KDs, however, is most likely due to pFD152 carriage and dCas9 expression. There was a reduction of PTS system, trehalose-specific IIBC component (TreB) and increase in xylose binding protein F (XylF) in *rlmJ* KD strains. The *rluD* KDs showed various proteins impacted including an increase in Phosphoadenosine 5’-phosphosulfate reductase H (CysH), Sodium/proline symporter (PutP), Tryptophan synthase beta chain (TrpB) and a reduction in Formate hydrogenlyase subunit 2 (HycB). For *rsmF*, a reduction in DNA-binding transcriptional activator (EvgA) was observed and conflicting changes (reduction in ATCC 11775 but increase in ATCC 8739) for Uncharacterized protein YqjD (YqjD). Only an increase in Oligopeptide transport system permease protein C (OppC) was seen for *rsmG*.

**Table 1.** List of proteins significantly changed due to gene knock down and detected in at least two strains.

| Gene targeted | Strain | Gene | Protein code<br>(UniParc) | Fold-change(log2) | -log10 (p-value) |
| --- | --- | --- | --- | --- | --- |
| *add* | 11775 | CitF | UPI00000DE86C | -2.93 | 3.47 |
|  | 8739 |  |  | -3.08 | 3.74 |
| *rlmF* | 11775 | CitF | UPI00000DE86C | -2.31 | 2.34 |
|  | 8739 |  |  | -3.51 | 3.45 |
| *rlmJ* | 11775 | CitF | UPI00000DE86C | -3.31 | 4.67 |
|  | 8739 |  |  | -3.25 | 5.60 |
|  | 11775 | TreB | UPI000000E8D4 | -2.25 | 2.79 |
|  | 25922 |  |  | -2.24 | 5.05 |
|  | 11775 | XylF | UPI0000DAD07E | 2.12 | 2.11 |
|  | 8739 |  |  | 2.07 | 3.58 |
| *rluD* | 11775 | CitF | UPI00000DE86C | -2.42 | 2.15 |
|  | 8739 |  |  | -2.70 | 2.60 |
|  | 25922 | CysH | UPI0000167DD2 | 2.43 | 4.62 |
|  | 8739 |  |  | 2.06 | 2.85 |
|  | 11775 | HycB | UPI0000D50BBA | -2.71 | 3.07 |
|  | 8739 |  |  | -2.59 | 2.78 |
|  | 11775 | PutP | UPI0000D507FA | 2.62 | 2.35 |
|  | 8739 |  |  | 2.56 | 3.26 |
|  | 25922 | TrpB | UPI00001639B2 | 2.07 | 5.36 |
|  | 8739 |  |  | 2.67 | 6.51 |
| *rsmF* | 25922 | EvgA | UPI000012A2A6 | -2.01 | 2.17 |
|  | 8739 |  |  | -2.25 | 3.22 |
|  | 11775 | YqjD | UPI0000047C93 | -4.45 | 2.01 |
|  | 8739 |  |  | 2.52 | 4.18 |
| *rsmG* | 11775 | OppC | UPI000000E133 | 2.18 | 2.07 |
|  | 8739 |  |  | 2.45 | 2.26 |

### Direct RNA Sequencing using Oxford Nanopore Technologies

ONT direct RNA sequencing was utilised to detect RNA modification sites. This dataset comprises of five known rRNA modification sites which include: *rlmF* (m6A, 23S rRNA, position 1618bp), *rlmJ* (m6A, 23S, 2030), *rluD* (Ψ, 23S, 1911/1915/1917), *rsmF* (m5C,16S, 1407) and *rsmG* (m7G, 16S, 527) (Supplementary Table S1) [16]. Two controls were included which were an initial isolate (pFD152 (-)) and pFD152 empty vector (pFD152 (+), dCas9 (+), no gRNA).

We utilised Nanocompore to identify putative RNA modification sites. Nanocompore does not require training of a model, is agnostic to the specific type of modification. Nanocompore does require two different conditions to compare, which is ideally suited to our experimental design where we have a KD state to compare to a control state.

Initially, flongle flow cells were used on total RNA extracts (∼90% rRNA) for ATCC 117755 to determine if known sites on rRNA could be detected and whether the CRISPRi model was successful. The majority of flongle generated reads (55.01%-73.72%) mapped to the rRNA reference (Figure 2B). Our results demonstrated that Nanocompore was able to identify known rRNA modification sites (Supplementary Figure S5).

**Figure 2.**
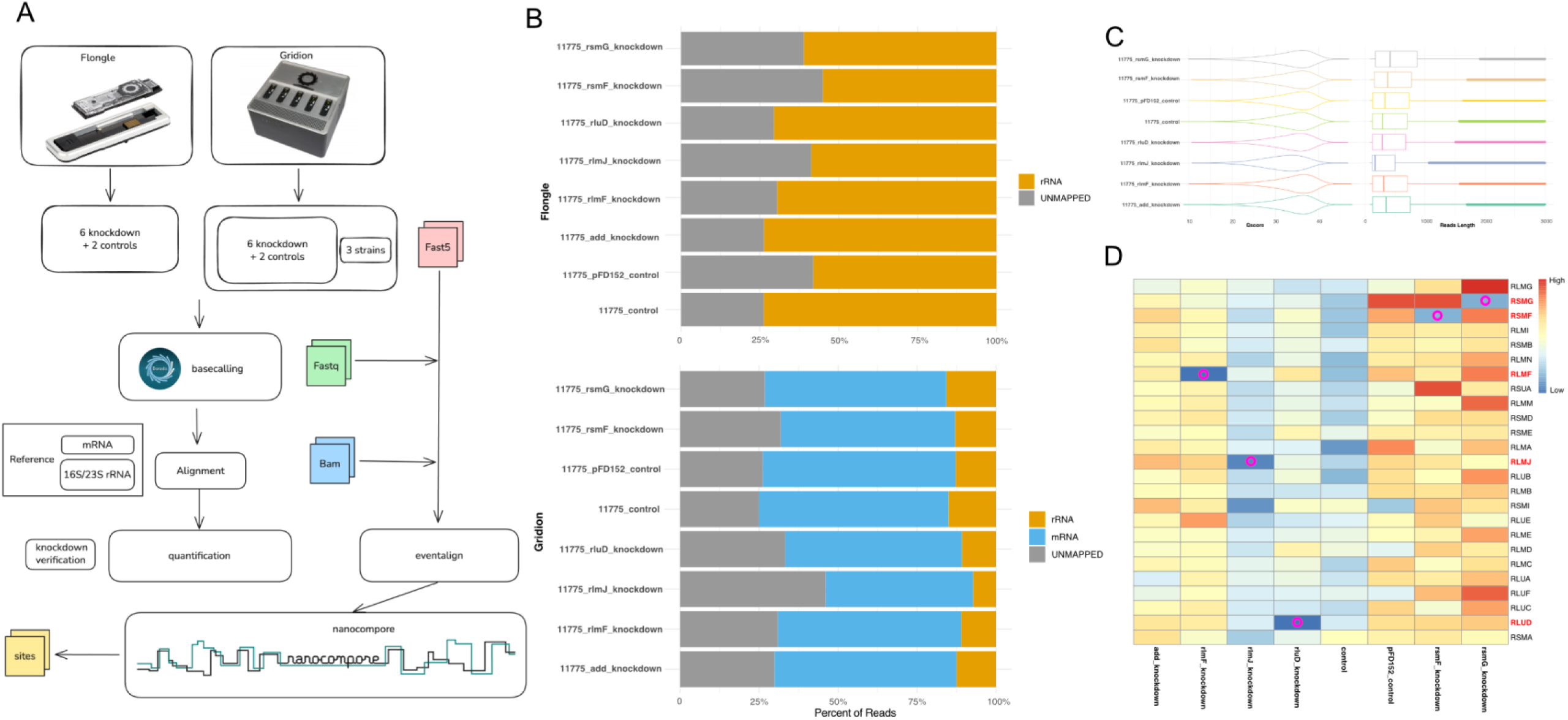
Nanopore direct RNA sequencing pipeline and analysis of ATCC 11775 *E. coli* controls and six CRISPRi samples. **A)** Workflow of sequencing and analysis. **B)** Proportion of reads mapping to reference (rRNA, mRNA) for flongle (rRNA) and GridION (mRNA enriched) sequencing runs. **C)** Quality of reads (Qscore) and read length for GridION (mRNA enriched) sequencing **D)** Heat map identifying repression of genes via CRISPRi detected using Salmon v1.10.1 for mRNA enriched sequencing. Circle indicates gene targeted via CRISPRi.

In order to generate enough coverage for detection of mRNA modifications, rRNA depletion was performed. mRNA enriched extracts were run on GridION flow cells (R9.4.1). For mRNA enriched ATCC 11775, only 7.22%-15.79% of reads mapped to rRNA whilst 46.76% - 61.05% mapped to mRNA (Figure 2B). Further, mRNA enriched samples for ATCC 8739 and ATCC 25922 predominantly (46.76%-75.69%) mapped to the mRNA reference (Supplementary Figure S6A). The ATCC 11775 GridION mRNA mapped reads had a median length of 165 – 419 and a Qscore of 32.1 to 34.7 (Figure 2C).

To confirm the KD efficiency, Salmon(v1.10.1) was implemented to quantify expression of genes of interest. We observed that the target KD genes expressed lower in the corresponding KD samples compared to controls (initial isolate, activated pFD152 (+) (dCas9 (+), no gRNA)) and samples with other KDs after in-sample and cross-samples normalization (Figure 2D). Similarly, for ATCC 8739 (Supplementary Figure S6C), the KD genes show lower expression in their corresponding KD sequencing samples. ATCC 25922 showed a minimal reduction in expression for KD genes of interest and was excluded from further analysis (Supplementary Figure S6B).

### Levels of rRNA site detection are dependent on gene targeted and modification type

Of the known rRNA targeted sites, the most significant signal via Nanocompore was with *rluD* (Ψ, 23S, 1911/1915/1917, reported motif: CGXAA, ACXAU, UAXAA) and *rsmG* (m7G, 16S, 527, reported motif: CCXCG) (Figure 3). The signal for *rlmF*, *rlmJ* and *rsmF* sites were difficult to detect and were challenging to identify above noise (Supplementary Figure S6). The three known Ψ modification sites for *rluD* (positioned in 23S rRNA stem-loop H69) (Figure 3A) could be detected for both flongle and GridION results against both controls (Figure 3B). This was apparent at sites between 1905-1920. Of note, the recognised nanocompore position number may have slight displacement (measured as a 5-mer with site + 3 nt) when comparing to the exact position in the genome. Volcano plots comparing KD and control reads identified significant changes at the known modification sites along with similar motifs detected (Figure 3C). Similarly, *rsmG* KDs showed significant signal at the known 16S rRNA site (Fig 3D-3F). The site with the most significance was at 521 for both flongle and GridION sequencing and previously reported motif CCXCG was detected.

**Figure 3.**
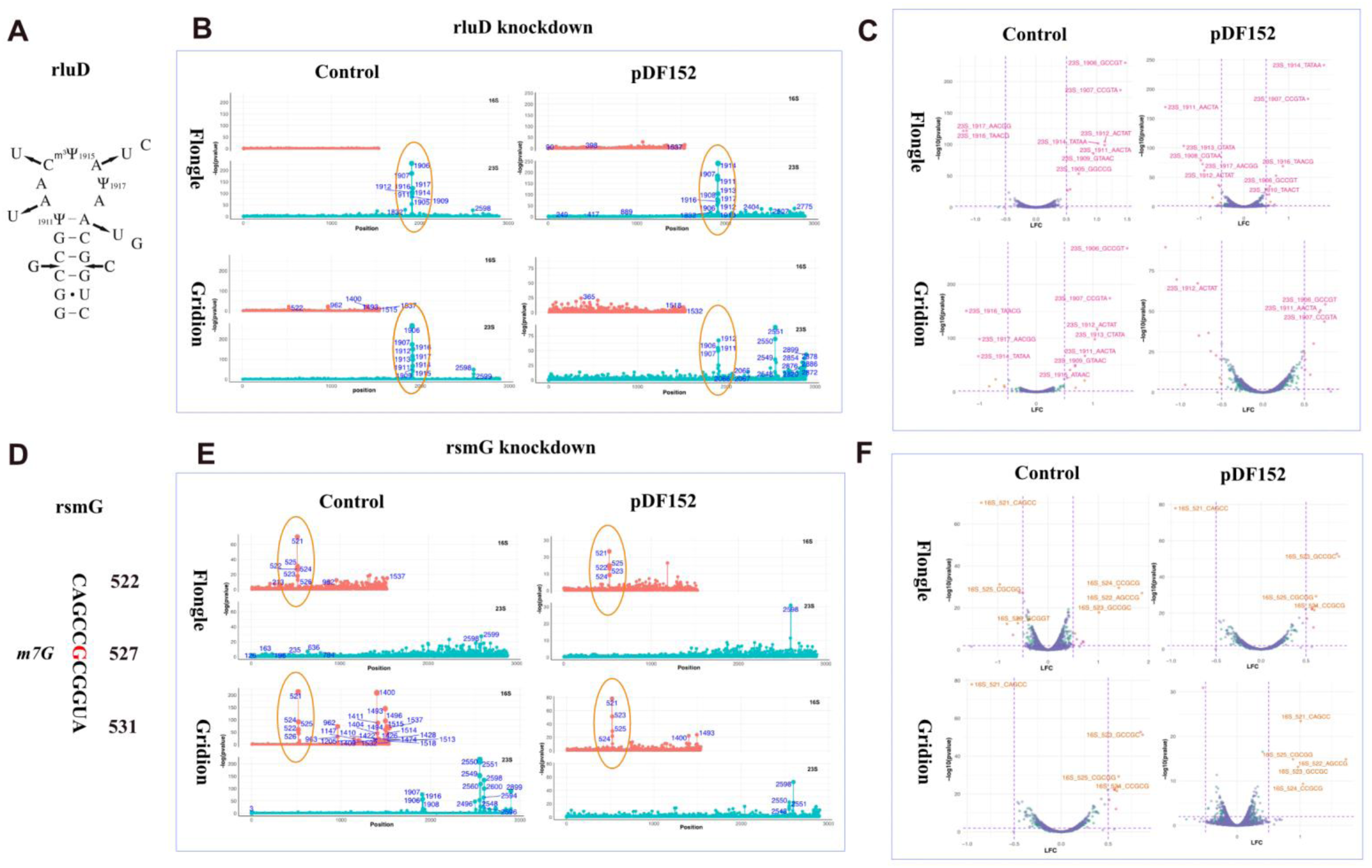
Nanocompore rRNA analysis of ATCC 11775 *E. coli* for *rluD* and *rsmG* CRISPRi compared against 2 controls. **A)** Modification (pseudouridine: Ψ) sites for *rluD* (23S, position 1911/1915/1917). **B)** Detection of *rluD* Ψ modification in flongle and GridION sequencing runs against 2 controls. **C)** Volcano plot detecting significant modification sites on rRNA and corresponding motif for *rluD*. **D)** Modification (7-methylguanosine: m7G) sites for *rsmG* (16S, position 527). **E)** Detection of *rluD* m7G modification in flongle and GridION sequencing runs against 2 controls. **F)** Volcano plot detecting significant modification sites on rRNA and corresponding motif for *rsmG*. Control contains no pFD152 plasmid (pFD152 (-)) and pFD152 control is pFD152 with activated dCas9 (pFD152 (+), dCas9 (+), no gRNA).

### ggcaller and mRNA modification site discovery

To unify the annotation, ggcaller (v1.3.0) was performed on ATCC 11775 and ATCC 8739 as these strains showed the most efficient KD (Figure 4A). For both ATCC strains, comparisons were run for each KD including KD vs control and KD vs pFD152. For each comparison, nanocompore compared the rate of modification from different conditions, KD and control, on each site on the mRNA the reads aligned to and provided a GMM p-value and fold change for each site. Following the suggestion from the nanocompore team, a threshold of p value < 0.01 was used and absolute value of fold change > 0.5 as cut off for significance. Motif analysis was performed using a logo plot [51] (Figure 4B) and summarised kmer patterns of significant sites in all pairs of comparison.

**Figure 4.**
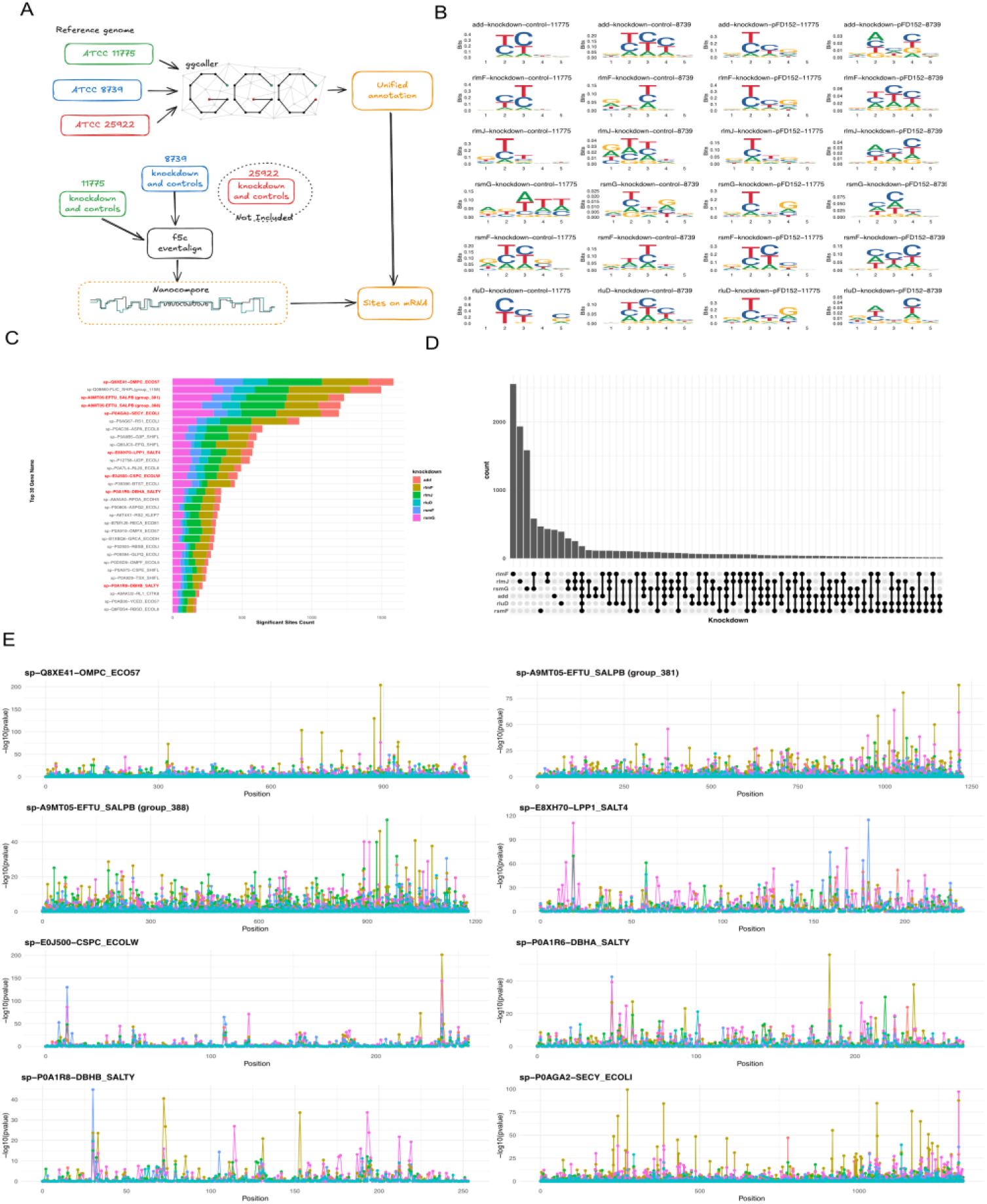
Nanocompore analysis of three ATCC *E. coli* strains to determine potential mRNA modifications. **A)** Workflow of analysis to construct a consensus reference and determine if the same mRNA modifications are detected between strains. **B)** Common mRNA motifs modified for six genes targeted against 2 controls in ATCC 11775 and ATCC 8739. **C)** Top 30 genes determined to harbour a modification site change due to CRISPRi. **D)** Number of genes with modification site changes similar across CRISPRi samples. **E)** Eight genes and modification signals corresponding to repression of genes targeted by CRISPRi. Control contains no pFD152 plasmid (pFD152 (-)) and pFD152 control is pFD152 with activated dCas9 (pFD152 (+), dCas9 (+), no gRNA).

Due to 4 pairs of comparison for each KD, Fisher’s method combined p values from different comparisons. A threshold p value of < 0.01 was applied to all sites on all mRNA. Sorting by number of significant sites, several genes which contained various significant modification sites were selected (Figure 4C). At the same time, we cross-checked the significant sites from different KDs to remove sites shared by different KDs, which are most likely artefacts. (Figure 4D). For each KD, the top 20 sites with smallest p values. (Supplementary Table S3) were selected, and there are several genes appearing in high frequently in the list. Using dot plots, we showed the significant sites distributed on the selected genes. (Figure 4E). Genes with significant modification sites include *ompC*, *lpp1, salPB, cspC, dbhA, dbhB* and *secY*. The motifs impacted on mRNA were further explored for *rluD* and *rsmG* KDs as they harboured the strongest signal shift on their known rRNA site (Table 2). For *rluD*, a significant change was found on the known motif (ACXAU) in gene EF-Tu (also known as *tuf1, tuf2*) position 891 and similar significant modification sites were detected in *lpp1* (position 44 (UAXUC)), 191 (UGXUA)), *g3p* or *gapA* (480 (GUXAU)) and *efg* or *fusA* (2019 (ACXAA))). For *rsmG*, a significant change was found on the known motif (CCXCG) in gene DBHA or *hupA* position 161 and similar significant modification sites were detected in *lpp1* (position 21 (GUXGG)), 126 (CAXCU)), *rl20* or *rplT* (227 (CAXCA)), *rs1* or *rpsA* (1516 (CCXGC)) and *ompC* (207 (CAXCU))).

**Table 2.** List of known rRNA motif sites and similar significant sites detected on mRNA.

| KD gene<sup>1</sup> | Mod | rRNA | Site | Known motif (X is base changed)<sup>2</sup> | mRNA<sup>3</sup> | Position | Motif<sup>4</sup> |
| --- | --- | --- | --- | --- | --- | --- | --- |
| *rluD* | Ψ | 23S | 1911<br>1915<br>1917 | CGXAA<br>ACXAU<br>UAXAA | A9MT05-EFTU_SALPB (group_388) | 891 | <u>ACCAU</u> |
|  |  |  |  |  | A0A4P7TN82-OMPF | 843 | GUUGC |
|  |  |  |  |  | E0J500-CSPC | 240 | GUCCG |
|  |  |  |  |  | E8XH70-LPP1 | 44 | <u>UACUC</u> |
|  |  |  |  |  |  | 58 | GUUGC |
|  |  |  |  |  |  | 120 | GUUGA |
|  |  |  |  |  |  | 168 | GCUGC |
|  |  |  |  |  |  | 191 | <u>UGCUA</u> |
|  |  |  |  |  |  | 212 | CAUGG |
|  |  |  |  |  | P0A1R6-DBHA | 87 | GCUGC |
|  |  |  |  |  | P0A9B5-G3P | 40 | GUCGC |
|  |  |  |  |  |  | 480 | <u>GUUAU</u> |
|  |  |  |  |  | P0AG67-RS1 | 1286 | CUCCC |
|  |  |  |  |  | Q83JC3-EFG | 2019 | <u>ACCAA</u> |
| *rsmG* | m7G | 16S | 527 | CCXCG | A9MT05-EFTU_SALPB (group_381) | 814 | CUGGC |
|  |  |  |  |  |  | 1069 | UCGAA |
|  |  |  |  |  |  | 1123 | UUGUU |
|  |  |  |  |  |  | 1163 | GCGUU |
|  |  |  |  |  | A9MT05-EFTU_SALPB (group_388) | 1006 | CUGAC |
|  |  |  |  |  | E8XH70-LPP1 | 21 | <u>CUGGG</u> |
|  |  |  |  |  |  | 126 | <u>CAGCU</u> |
|  |  |  |  |  |  | 141 | GUGAA |
|  |  |  |  |  |  | 204 | CUGGA |
|  |  |  |  |  |  | 213 | AUGGC |
|  |  |  |  |  | P0A1R6-DBHA | 161 | **<u>CCGCG</u>** |
|  |  |  |  |  | P0A1R8-DBHB | 114 | CUGGU |
|  |  |  |  |  | P0A7L4-RL20 | 227 | <u>CAGCA</u> |
|  |  |  |  |  |  | 259 | CUGUU |
|  |  |  |  |  | P0A975-CSPE | 116 | UGGUU |
|  |  |  |  |  | P0AG67-RS1 | 1516 | <u>CCGGC</u> |
|  |  |  |  |  | Q8XE41-OMPC | 207 | <u>CAGCU</u> |

<sup>1</sup>KD genes *rluD* and *rsmG* only investigated further due to strong signal on known rRNA site.
<sup>2</sup>Motifs as per: Fleming AM, Bommiseti P, Xiao S, Bandarian V, Burrows CJ. Direct Nanopore Sequencing for the 17 RNA Modification Types in 36 Locations in the *E. coli* Ribosome Enables Monitoring of Stress-Dependent Changes. ACS Chem Biol. 2023 18:2211-2223. doi: 10.1021/acscchembio.3c00166.
<sup>3</sup>Gene name as per output from ggCaller. mRNA sites reported required a significant signal for both ATCC 11775 and ATCC 8739.
<sup>4</sup>Motifs selected required X to be U or C for *rluD* (Ψ can be basecalled as a C: Fleming AM, Mathewson NJ, Howpay Manage SA, Burrows CJ. Nanopore Dwell Time Analysis Permits Sequencing and Conformational Assignment of Pseudouridine in SARS-CoV-2. ACS Cent Sci. 2021 7:1707-1717. doi: 10.1021/acscentsci.1c00788.) and *rsmG* as a G. represents exact match and indicates a 1-2 nt difference from known motif.

## DISCUSSION

CRISPRi is a newly developed genetic engineering approach which has shown promise to KD genes in bacteria [29–32]. In this study, we selected CRISPRi rather than CRISPR as there can be high toxicity associated with cutting the genome. We further selected an optimised CRISPRi system which has been optimised for *E. coli* and harbours an inducible dCas9 (via aTc) as constant expression of dCas9 is toxic [29]. Whilst CRISPRi holds promise, several limitations were observed in this study. Firstly, finding PAM sites close to the start site of the gene were challenging. Secondly, bacterial genes are tightly clustered and commonly regulated via operons in which several genes are transcribed on the same RNA molecule. Hence, halting the transcription at the start of an operon can inhibit several genes and discerning the impact of one gene of interest is not possible. For *E. coli*, a curated database exists (RegulonDB) where operons can easily be identified, however, this is lacking for other bacterial species. Thirdly, expressing the pFD152 plasmid (9 kb) can result in growth defects and compete with the accessory genome as evident with ATCC 25922 and ATCC BAA-2452. There are alternative models for CRISPRi including integrating dCas9 into the chromosome with a strong promotor to enable sufficient dCas9 expression and repression [34]. Despite these shortcomings, we observed a substantial reduction in expression of genes targeted via CRISPRi (∼>80%) in ATCC 11775 and ATCC 8739.

Activation of dCas9 in ATCC 11775 caused an overall growth delay whilst ATCC 8739 revealed no impact on growth when dCas9 was induced. This may be attributed to ATCC 11775 having an additional 131 kb plasmid along with pFD152. When comparing with controls, a growth delay was observed in *rluD* and *rsmF* in ATCC 11775 and *rlmF*, *rluD* and *rsmF* KDs in ATCC 8739. *Pletnev* et al [52] explored the growth of *rlmF*, *rlmJ*, *rsmF* and *rsmG* KO *E. coli* (strain BW25113) and identified no significant change. A prior study by Gutierrez *et al*. identified that deleting *rsmF* can reduce growth in *Pseudomonas aeruginosa* [53] however, the contrary was observed for *E. coli* by Lioy *et al.* [54]. In our work, we note a significant growth reduction associated with *rsmF* however, experimental design may attribute to this (number of cells seeded per well, time between OD600 measurements) and may detect more subtle growth delays. Repression of *rluD* in both strains also caused growth delays. O’Conner *et al.* observed conflicting results for growth and highlight this result depends on the genetic background of the *E. coli* strain (e.g. mutations in *prfB* and *prfC* genes) [55]. We identified a growth delay in *rlmF* KD in ATCC 8739 and this has previously been reported in *rlmF* knock out *E. coli* [55]. Sergiev *et al.* also find a minor growth delay in *rlmJ* KOs and whilst we did not observe a significant delay for *rlmJ* KDs, on average there was a growth reduction. Overall, experimental design and differing *E. coli* strains used may explain this high variability in reporting growth delays.

RNA modifications can influence the proteome especially those present on rRNA which are responsible for protein synthesis. These rRNA modifications can fine-tune translational efficiency and enable survival under certain environmental conditions (cold temperatures, antimicrobials) in bacteria [52]. As we target 5 rRNA modification genes in this study, we sought to determine the influence on the proteome. Pletnev *et al* [52] interrogated the proteome of *rsmF* KO *E. coli* and identified numerous protein changes. We compared the significant protein changes found for *rsmF* KD in ATCC 11775 and ATCC 8739. The exact proteins impacted were not observed; however, similar gene families such as PspE (decrease) and YbeD (decrease) (Pletnev *et al* [52]) and PspB (increase) and YbeY (decrease) in ATCC 11775 in our study. Differing *E. coli* strains, protein extraction protocol and mass spectrometer were used in this prior study. Limited studies have been conducted on the link between rRNA modification loss and proteome in bacteria. We are the first to report these protein changes linked to rRNA modification loss in ATCC 8739 and 11775: XylF (*rlmJ* KD), *HycB* and *PutP* (*rluD* KD) and *OppC* (rsmG KD). Overall, only minor protein changes were observed in our study. It has previously been noted that particular conditions (such as antimicrobial stress, temperature changes) may be required to determine the functional role of these rRNA modifications [16, 25, 52].

Direct RNA sequencing is a promising approach to rapidly detect the epitranscriptome but has rarely been applied to bacterial RNA. Fleming *et al* [16] provided a comprehensive overview for detecting all *E. coli* (strain K12 DH5α) modified rRNA sites using direct RNA sequencing. This study utilised the same chemistry (SQK-RNA002, R9.4 flow cells) as our study but differed by the growth phase (stationary rather than log phase). This work detected RNA modifications using differing tools including Nanopore-Psu and ELIGOS2 to quantify base call data, Nanopolish, Nanocompore and Tombo to determine ionic current and helicase dwell time. Fleming *et al.* highlighted the challenges to detect m5C modifications for all 3 sites on *E. coli* rRNA which was similar to our work (*rsmF*, m5C modification). The detection of m6A (*rlmF*, 23S rRNA, 1618bp) was very weak when using tools based on base calling and current with only helicase dwell time as an indicator. The m6A modification via *rlmJ* (23S rRNA, 2030bp) signal was stronger with evidence of this site modification via base calling and dwell time tools. Other modifications including Ψ and m7G had stronger signals. Sites modified by *rluD* (23S rRNA, Ψ1911/1915/1917) could be determined via basecalling, current and dwell time however, current provided ambiguous sites due to the close proximity of modifications and dwell time missed site 1917. For m7G (*rsmG*, 16S rRNA, 527bp), base call and current could detect this site but not dwell time which align with our results. This work also explored whether metabolic stress (minimal media) and temperature (20°C) can alter the RNA modification detection. Due to the weak signal, sites for all m5C sites and m6A (site 1618, *rlmF*) were not measured. Under metabolic and temperature stress, m6A addition via *rlmJ* (23S rRNA, 2030bp) significantly decreased (p<0.05). No change for m7G position 527 (16S rRNA, *rsmG*) was observed under metabolic and temperature stress. No change was detected for Ψ at positions 1911/1915/1917 (*rluD*, 23S rRNA) under metabolic stress however, interestingly, a significant decrease (p<0.05) was evident under temperature stress in which Ψ is known to be a heat stabilising RNA modification. Antimicrobial stress can also influence the detection of RNA modifications via direct RNA sequencing as highlighted by Delgado-Tejedor *et al* [57]. Hence, this highlights RNA modification abundance and detection for direct RNA sequencing is strongly influenced by growth conditions.

Recently, two key studies have performed an in-depth investigation of direct RNA sequencing and further benchmarked the RNA modification detection potential on bacterial transcripts. Tan *et al* [58] performed direct RNA sequencing on *E. coli* (K-12) and *S. aureus* (Wichita) using the SQK-RNA002 kit and *in vitro* transcription as a negative control (removing RNA modifications). As m6A is a key modification and many tools have been trained on this change, their study further used immunoprecipitation (MeRIP-Seq) to enrich for these sites on mRNA. Whilst this study used various tools to detect modification sites, nanocompore was able to detect various m6A sites although, may have missed several sites compared to DRUMMER [59] and Differr [60]. This work highlights that the commonly known motif for m6A modifications, UGCCAG [17], was not the highest hit in the dataset rather, GACGCMAG (M: C/A), in *E. coli*. Several high confidence m6A sites on mRNA were also identified. When comparing these sites to our m6A sites (*rlmF*, *rlmJ*), we also detected potential m6A changes on *ompC* and *secY*. Tan *et al* noted a strong m6A signal on *ompC* at position at 167 bp. We further investigated this site and observed a significant change at position 167 for *rlmJ* KD however, the motif differed (known motif: UGXAG, detected: CUX(CA or UG)). For *secY*, position 23 had a strong signal however, we did not see a significant signal at this position. This work was further expanded by Guo *et al* [61] who used the new direct RNA sequencing chemistry (SQK-RNA004, FLO-MIN004RA flow cells) which has higher median accuracy (≥97.5% across modified and unmodified (IVT) datasets) and a 4-fold increase in data yield. Guo *et al* enhance the detection of bacterial RNA modifications with direct RNA sequencing and develop a new tool, nanoSundial (available for kit RNA004 only). This study discovers 190 sites stably modified on highly expressed mRNA in *E. coli* however, the exact modification type is not known. Overall, the release of the newer version of direct RNA sequencing is promising for rapidly detecting RNA modifications. However, further fine-tuning is required for mRNA modification detection across differing bacterial species including modification type and dynamics under certain growth conditions.

## ACKNOWLEDGEMENTS

The authors would like to acknowledge Dr. Dianna M. Hocking for her microbiology expertise and input for CRISPRi experiments. We acknowledge Dr. Jessica M. Lewis for initial assistance with CRISPRi and proteome experimental design. The authors further acknowledge Kevin Chen and Megan Spralja for initial assistance with CRISPRi experiments. We thank the Melbourne Mass Spectrometry and Proteomics Facility of The Bio21 Molecular Science and Biotechnology Institute for access to MS instrumentation. We would like to acknowledge the Media Preparation Unit (University of Melbourne) for their quick turnaround for custom media and agar plates.

## AUTHOR CONTRIBUTIONS

M.E.P. and L.J.M.C conceived the study and obtained research funding. M.E.P, N.E.S and L.J.M.C. designed the experiments. M.E.P, A.N.T.N. and L.J. performed the experiments. M.E.P, J. Z., A.N.T.N., M.B.H., L.A.F, N.E.S and L.J.M.C. analysed the data and prepared figures. M.E.P, J. Z., A.N.T.N., M.B. H., L. J., L.A.F., G. S. A. M., N.E.S and L.J.M.C interpreted results. M.E.P. wrote the first draft of the paper. All authors edited and reviewed the paper.

## ETHICS

Generation of CRISPRi *E. coli* were approved under NLRD IBC reference number: 2020.066 (University of Melbourne).

