## Supplementary Figures and Tables for "Interrogating the *Escherichia coli* epitranscriptome via CRISPR interference and Nanopore native RNA sequencing"

### SUPPLEMENTARY MATERIAL

Miranda E. Pitt[1,2*,§], Jianshu Zhang[1§], An N. T. Nguyen[1], Michael B. Hall[1], Leila Jebeli[3], Leo A. Featherstone[2,4], Garry S.A. Myers[1], Nichollas E. Scott[1] and Lachlan J.M. Coin[1*]

[1]Australian Institute for Microbiology and Infection, University of Technology Sydney, New South Wales, Australia

[2]Peter Doherty Institute, University of Melbourne, Victoria, Australia

[3]Enteric Diseases Group, Murdoch Children's Research Institute, Victoria, Australia

[4]The Kirby Institute, University of New South Wales, Sydney, Australia

§ These authors contributed equally

**TABLES**

**Table S1**. Genes targeted via CRISPRi, PAM site positions, location in genome and score for off-target sites

| Gene | Modification, RNA type, position (bp) | gRNA | Sequence (Forward)[#] | Gene length (bp) | Gene position[‡] | PAM start position (bp)[^] | Score (minimal off-targets)[§] |
|---|---|---|---|---|---|---|---|
| *add* | - | 1[*] | TAGTCAGGAAGCGAGATATTATAC | 1002 | 2383714 - 2384715 | +89 | 0.81 |
| | | 2 | TAGTTCATGGTCGCACTCTTTTTC | | | +110 | 0.29 |
| *rlmF* | m6A, 23S rRNA, 1618 | 1[*] | TAGTAACGCGACAGGGTAAATATC | 927 | 3215455 - 3216381 | -59 | 0.71 |
| | | 2 | TAGTCATTTCTAAAAACCTTCACA | | | -92 | 0.64 |
| *rlmJ* | m6A, 23S rRNA, 2030 | 1[*] | TAGTGTGTTCCGGTAAGTAAAAAT | 843 | 237266 - 238108 | -29 | 1.47 |
| | | 2 | TAGTGAGCATGGGTAAAGGTGTTC | | | -15 | 0.79 |
| *rluD* | Ψ, 23S rRNA, 1911/1915/1917 | 1[*] | TAGTGCGTTGACCGAGTTGGTTTT | 981 | 1280270 - 1281250 | +38 | 0.99 |
| | | 2 | TAGTATATAGTGTGCTATTGTAGC | | | -70 | 0.84 |
| *rsmF* | m5C, 16S rRNA, 1407 | 1[*] | TAGTCGGGAAATAAACGGTGTGTT | 1440 | 2178852 - 2180291 | +4 | 0.86 |
| | | 2 | TAGTGCAGTTTAGCATAAACGCTC | | | -54 | 0.70 |
| *rsmG* | m7G, 16S rRNA, 527 | 1[*] | TAGTAGGCAATAAGCTGGTTTTTC | 624 | 4864256 - 4864879 | +56 | 0.92 |
| | | 2 | TAGTAAGCTGGTTTTTCTGGTGAT | | | +49 | 0.52 |

[*]gRNA construct selected for subsequent experiments determined via a combination of best gene position, fewer off-target sites and higher knock down efficiency based on qRT-PCR results.

[#]Reverse gRNA is 5’-AAAC following the reverse compliment of forward sequence. Flanking for insertion via BsaI restriction site.

[‡]Gene positions as per ATCC11775 genome (NZ_CP033092.2) available via NCBI. Represented as start-end position in genome.

[^]PAM (5′-NGG-3′) binding site location represented as before (-) or after (+) start position in gene.

[§]Off target score determined using the crispr-browser.pasteur portal [1]. Higher score indicates fewer off target sites for dCas9 to bind to the genome.

**Table S2.** Oligos used for qRT-PCR to determine gene knock down efficiency and PCR to confirm gRNA in plasmid

| Gene | Sequence (Forward) | Sequence (Reverse) | Alternative oligo |
|---|---|---|---|
| **qRT-PCR** | | | |
| *add* | CTGGGGCGTTAAAGTTCTCG | TACGGGCTGCATCTTCAATG | GACGGGCTGCATCTTCAATG (Reverse)[#,‡] |
| *rlmF* | GAGTGGGGCGATTTTTAACG | ATGGCGGGTTACACAAGGTC | |
| *rlmJ* | CCGCCGTGGTTTAATCCTTA | CCAGTGGCGAAACGTTTGTA | |
| *rluD* | GAAGAAGCGCGTTTTGAACC | ACCAGGTCGCGTGGTTTATT | ACCAGGTCGCGCGGTTTATT (Reverse)[#,‡] |
| *rsmF* | GTACCGCCGAGCATTTAAGC | GGTGCATTACCGTCAGCAAA | GGTGCATTATCGTCAGCAAA (Reverse)[‡] |
| *rsmG* | TCCTTCGTCAGGTGCAACAT | CGGCTCTGACGGAAACTCTT | CGGCTCTGAAGGAAACTCTT (Reverse)[#,‡] |
| *rpoS*[#] | TTGCTGGACCTGATCGAAGA | GAGAAGCGGAAACCACGTTC | TTGCTGGACCTTATCGAAGA (Forward)[#,‡] |
| *rssA*[#] | AGGACTAATGGCACCCGTTG | ACGCGTGAGGGAAATAGGAA | AGGACTCATGGCACCTGTTG (Forward)[#,‡] |
| **PCR** | | | |
| pFD152 plasmid[^] | GTCTTTATGAAACACGCATTGA | GGGAATGAGAATAGTGAATGGACC | |

[#]Housekeeping genes selected based on prior literature [2,3].

[#]qRT-PCR alternative primer required for isolate ATCC8739.

[‡]qRT-PCR alternative primer required for isolate ATCCBAA2452.

[^]Oligos used to PCR flanking regions of pFD152 with gRNA and sequence checked via Sanger sequencing. To check gRNA was present, pFD152 Forward underwent PCR with gRNA Reverse oligo.

**Table S3**: Gene positions with unified ggcaller annotation

| ggcaller_ID[*] | Gene | Alt | 11775 | 8739 | 25722 |
|---|---|---|---|---|---|
| sp-E8XH70-LPP1_SALT4 | LPP1 | | 477018-477254 | 2159408-2159644 | 2381476-2381712 |
| sp-Q8XE41-OMPC_ECO57 | OMPC | | 4761941-4763032 | 1687186-1688313 | 1582947-1584077 |
| sp-E0J500-CSPC_ECOLW | CSPC | | 336248-336508 | 2008789-2009049 | 2236106-2236366 |
| sp-A9MT05-EFTU_SALPB (group_388) | EFTU | TUF1, TUF2 | 2726375-2727559 | 4456195-4457379 | 4808415-4809599 |
| sp-A9MT05-EFTU_SALPB (group_381) | EFTU | TUF1, TUF2 | 3464087-3465316 | 423510-424739 | 433585-434814 |
| sp-P0A1R6-DBHA_SALTY | DBHA | HUPA | 2703993-2704265 | 4433689-4433961 | 4786031-4786303 |
| sp-P0A1R8-DBHB_SALTY | DBHB | HUPB | 1688907-1689107 | 3498197-3498454 | 3757228-3757485 |
| SP−P0AGA2−SECY_ECOLI | SECY | | 3491391-3492722 | 450787-452118 | 460874-462205 |

[*]ggcaller ID includes uniprot protein reference in description

[#]Alternative name for gene

**FIGURES**

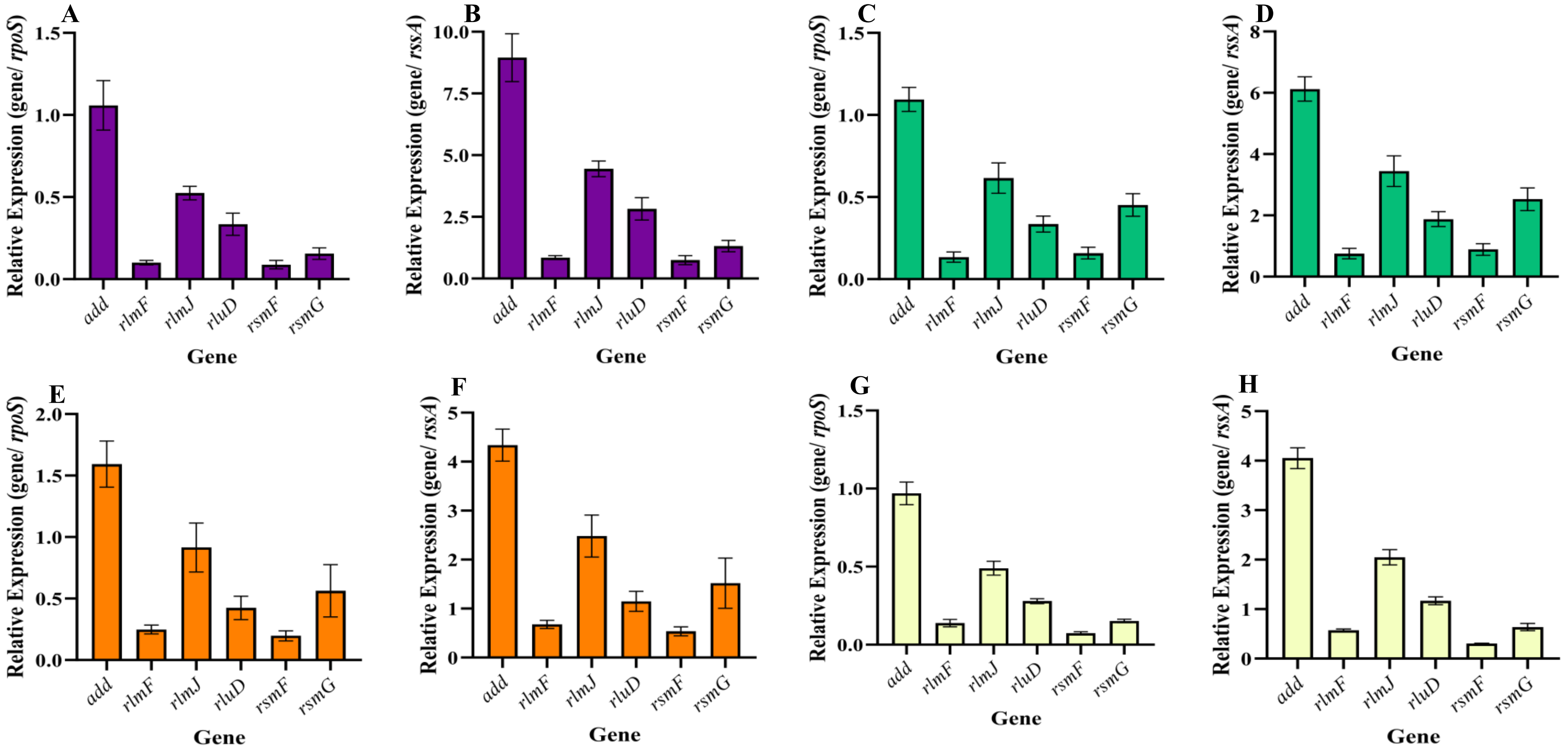


**Figure S1.** Expression of six genes across four *E. coli* strains and normalised to two housekeeping genes, *rpoS* and *rssA*. **A)** ATCC 11775 *rpoS*. **B)** ATCC 11775 *rssA*. **C)** ATCC 25922 *rpoS*. **D)** ATCC 25922 *rssA*. **E)** ATCC 8739 *rpoS*. **F)** ATCC 8739 *rssA*. **G)** ATCC BAA-2452 *rpoS*. **H)** ATCC BAA-2452 *rssA*. Bars represented as average (n=3) ±standard deviation.

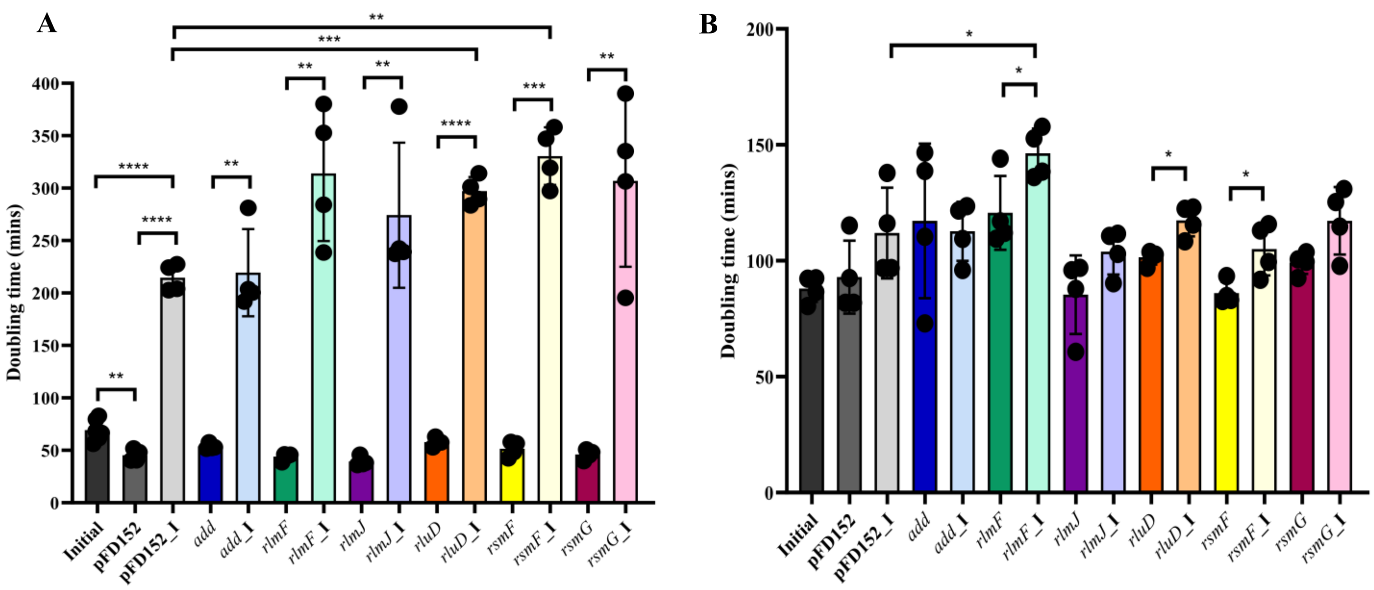


**Figure S2.** Growth rate comparison between controls (Initial) and KD (pFD152) for two *E. coli* strains. **A**) ATCC 11775. **B**) ATCC 8739. Mean±SD, n≥4 biological replicates, "_I" is CRISPRi - activation of dCas9+gRNA. Three controls were included for each strain: "Initial" (first column) represents ATCC strain with no pFD152 plasmid, "pFD152" (second column) is pFD152 within the strain but not activated (spectinomycin selection only, no gRNA) and "pFD152_I" is pFD152 within the strain and activated (spectinomycin selection and anhydrotetracycline supplementation, no gRNA, dCas9 expression). Significance represented as *p<0.05, **p<0.01, ***p<0.001, ****p<0.0001.

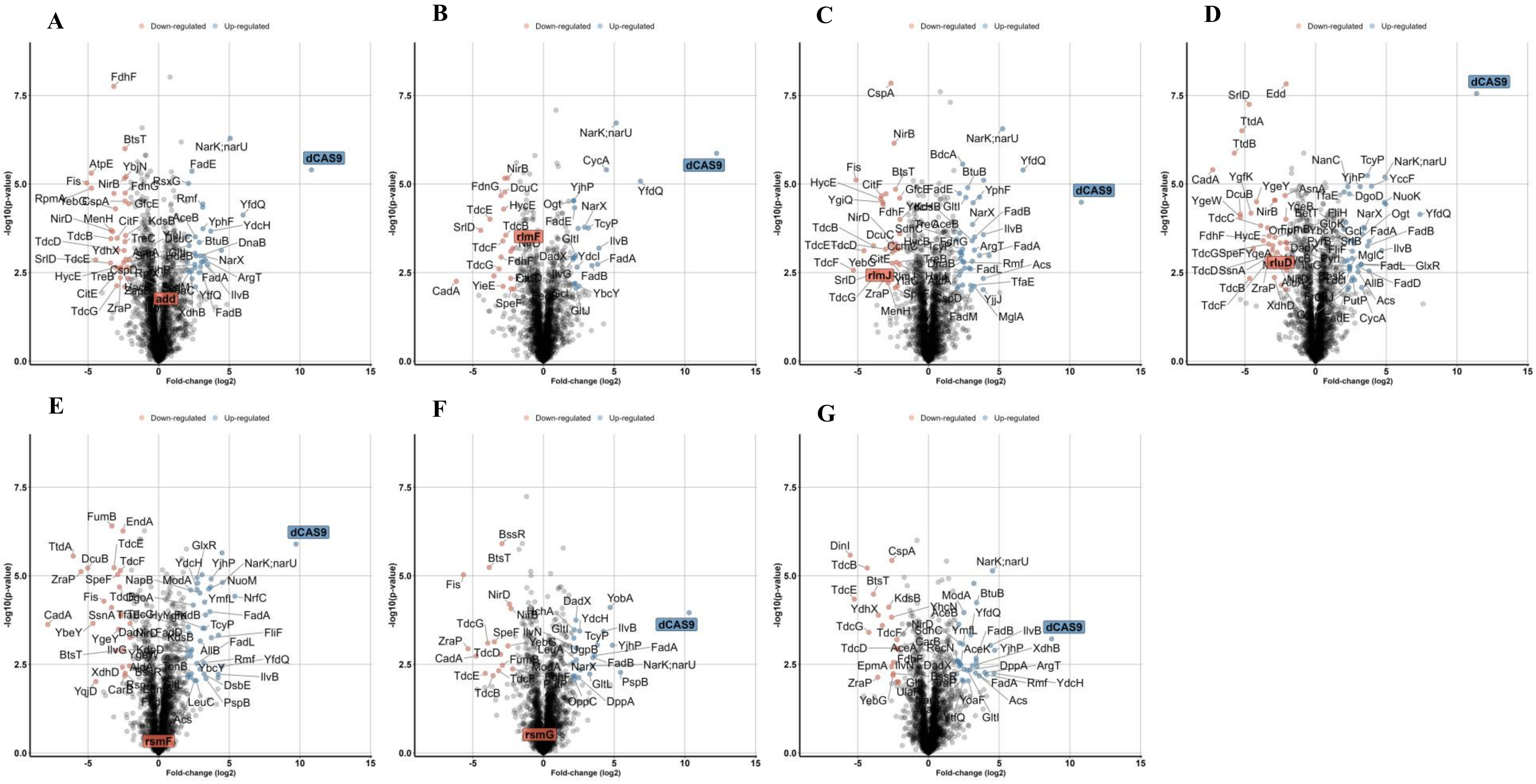


**Figure S3.** ATCC 11775 protein changes when comparing control (pFD152 plasmid carriage only) and knock down (expression of dCas9 and gRNA) samples. **A**) *add*. **B**) *rlmF*. **C**) *rlmJ*. **D**) *rluD*. **E**) *rsmF*. **F**) *rsmG*. **G**) pFD152. Protein levels of genes KD and dCas9 shaded. Down- (orange) and up-regulation (blue) defined as fold-change(log2) >2 or <-2 and -log10 (p-value) >2. n = 4 biological replicates.

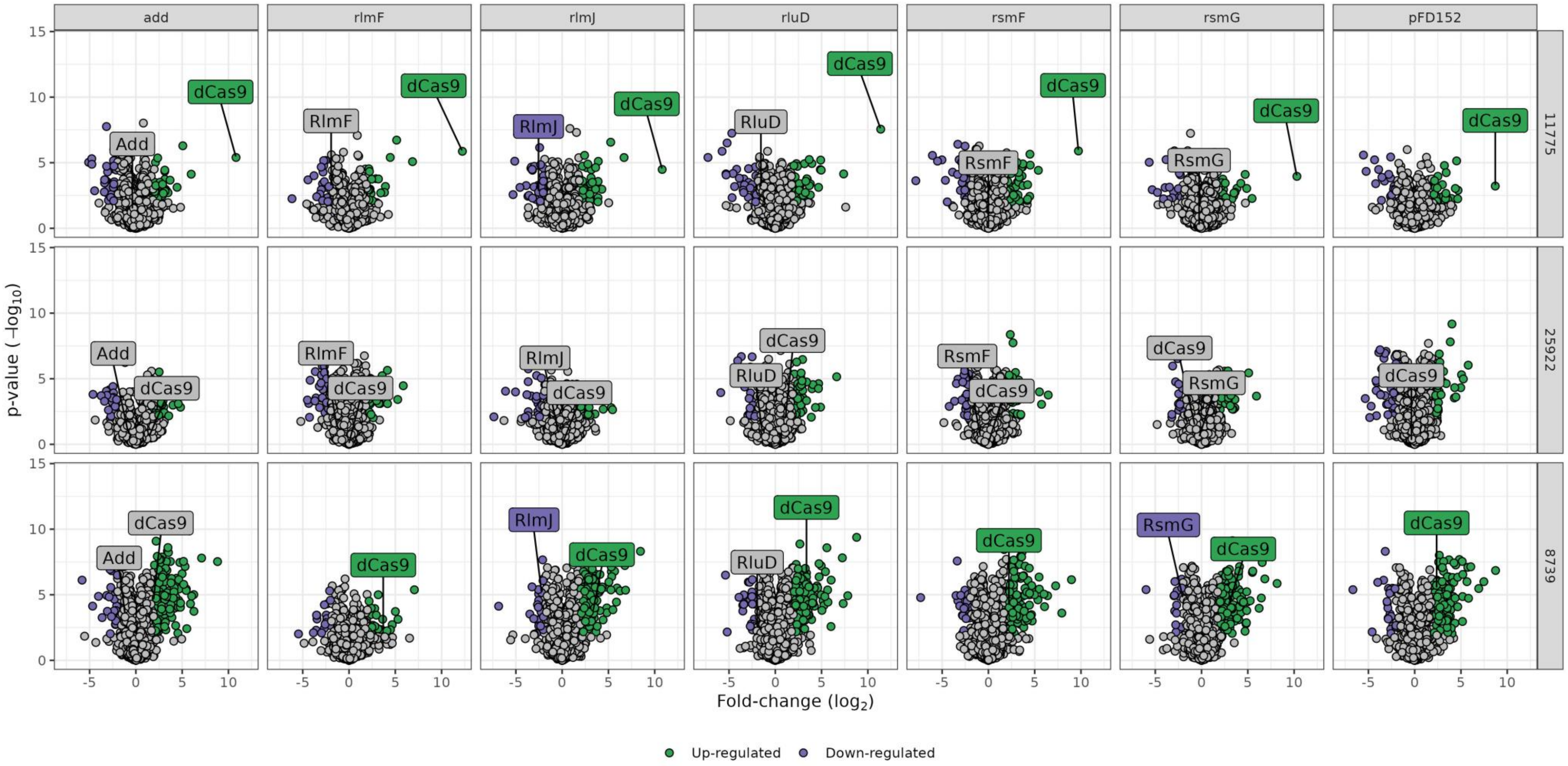


**Figure S4.** Protein changes in control with pFD152 plasmid not induced to express dCas9 and gRNA vs pFD152 knock down in three ATCC *E. coli* strains. Down- (purple) and up-regulation (green) defined as fold-change(log2) >2 or <-2 and -log10 (p-value) >2. n=4 biological replicates

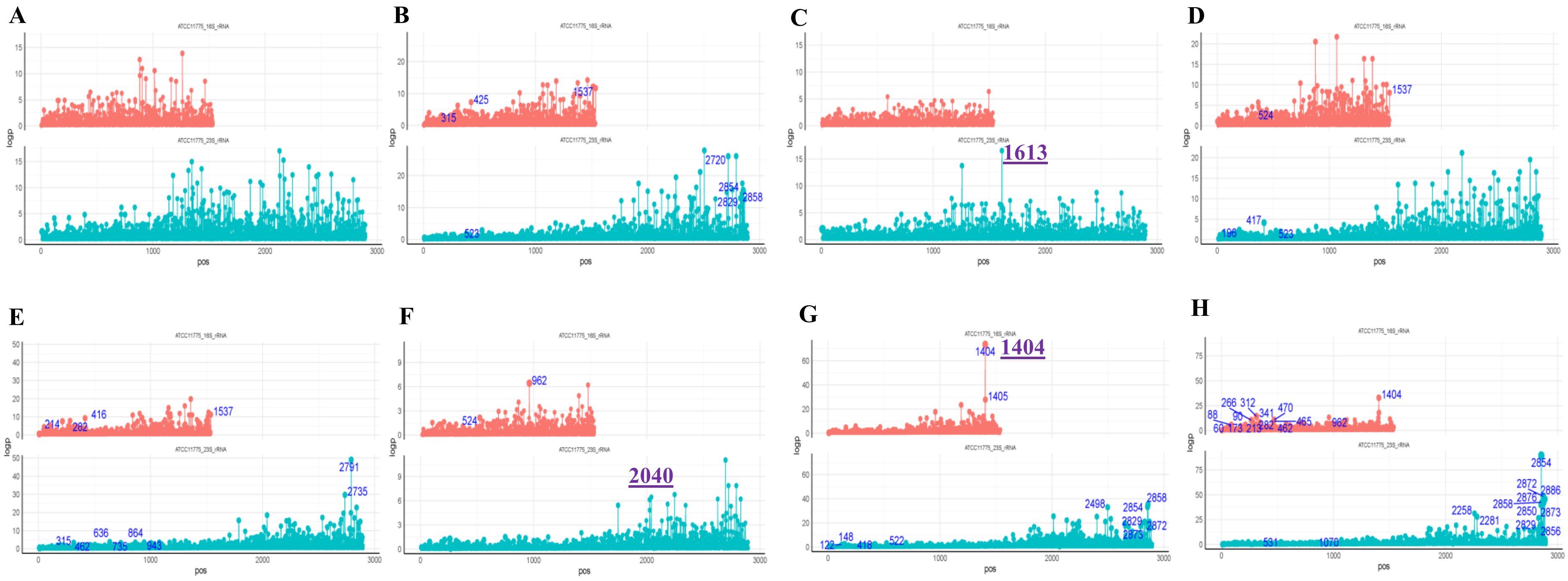


**Figure S5.** Nanocompore detection of known rRNA modification sites using flongle (total RNA) data. **A)** *add* CRISPRi vs initial control. **B)** *add* CRISPRi vs pFD152 control. **C)** *rlmF* CRISPRi vs initial control. **D)** *rlmF* CRISPRi vs pFD152 control. **E)** *rlmJ* CRISPRi vs initial control. **F)** *rlmJ* CRISPRi vs pFD152 control. **G)** *rsmF* CRISPRi vs initial control. **H)** *rsmF* CRISPRi vs pFD152 control. Initial control contains no pFD152 plasmid and pFD152 control is pFD152 with activated dCas9 (supplemented with spectinomycin and anhydrotetracycline). **Purple bold underlined** position indicates detection of known site.

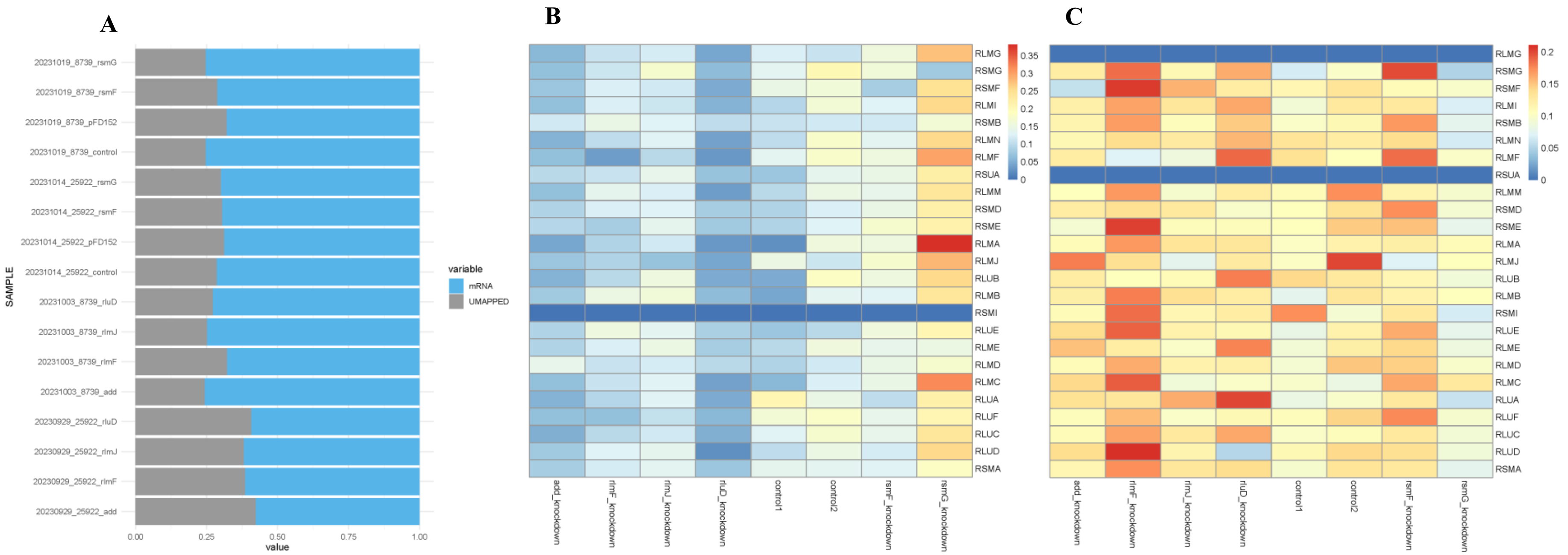


**Figure S6.** Nanopore direct RNA sequencing of ATCC 25922 and ATCC 8739 *E. coli* controls and six CRISPRi samples. **A)** Proportion of reads mapping to reference (mRNA) for GridION (mRNA enriched) sequencing runs **B)** ATCC 25922 heat map identifying repression of genes via CRISPRi detected using Salmon v1.10.1 for mRNA enriched sequencing. **C)** ATCC 8739 heat map identifying repression of genes via CRISPRi detected using Salmon v1.10.1 for mRNA enriched sequencing. Control 1 is initial control which contains no pFD152 plasmid (pFD152 (-)) and control 2 is pFD152 with activated dCas9 (pFD152 (+), dCas9 (+), no gRNA).
